# Computational ligands to VKORC1s and CYPs. Could they predict new anticoagulant rodenticides?

**DOI:** 10.1101/2021.01.22.426921

**Authors:** A Bermejo-Nogales, J.M. Navas, J Coll

## Abstract

Anticoagulant-resistance in rodents and anticoagulant off-target effects are some of the world-wide problems of increasing concern. To search for new anticoagulant rodenticide candidates we have computationally explored some of the rat genes previously implicated in resistance to actual anticoagulants. In particular, we searched among hundreds of anticoagulant-similar chemotypes those binding rat wild-type VKORC1 (the best-known anticoagulant target, a Vitamin K-recycling enzyme), VKORC1L1 (a VKORC1-related enzyme), Cytochrome P450 CYP enzymes (some of the most important enzymes implicated in detoxification) and anticoagulant-resistant VKORC1-mutants (to minimize propensity to resistance). Results predicted new VKORC1 leads with binding-scores in the low nM range (high binding-affinities) predicting hydroxycoumarin- and naphtoquinone-like chemotypes. We then selected top-leads with additional high binding-scores to more than three anticoagulant-related CYPs, suggesting minimal detoxification rates and therefore maximal anticoagulation expectatives. A downsized list of top top-leads maintaining VKORC1 low-binding scores to anticoagulant resistant mutants, was finally proposed for experimental validation. The combination of different rat targets for computational studies, could be used to search for unrelated chemotypes, for reduction of off-target environmental anticoagulant impacts, and/or as new tools to explore anticoagulant molecular mechanisms.

## Introduction

Anticoagulant rodenticides have been used during the last decades to control populations of the wild rat, *Rattus norvegicus* (Berk.). It was from the 50’s that rat genetic resistances to anticoagulants were increasingly detected^1, 2^. Many field surveys showed resistance to rodenticide anticoagulants not only to those of the first-generation, such as warfarin (58 mg/Kg LD^50^) or coumatetralyl (16.5 mg/Kg), but also to the more potent second-generation, such as difenacoum (1.8 mg/Kg), bromadiolone (1.1 mg/Kg), difethialone (0.56 mg/Kg), flocoumafen (0.46 mg/Kg) or brodifacoum (0.26 mg/Kg)^3, 4^. Despite being more potent, the second generation anticoagulants also showed a practical requirement for delayed mortality to reduce rats of “ learning” the association of feed on poisoned baits with faster deaths. On the other hand, anticoagulant chemical residues have been increasingly detected on the tissues of several off-target wildlife species ^1, 3, 5-7^, generating additional concerns which also include humans. To mitigate such environmental risks, in several countries the more-potent anticoagulant rodenticides are only allowed to use in or nearby invaded buildings and only after there is proof of resistance to less-potent anticoagulants. Although both combination with other drugs (for instance, coumatetralyl and cholecalciferol) ^8^ and proposed novel brodifacoum-like compounds computationally predicted with higher binding affinities to the main target enzyme ^9^ have been previously reported, there is an urgent need for new anticoagulant rodenticides and/or combinations which could maximize on-target and minimize off-target effects. Alternatively, other rodent-specific targets may also be investigated, but those are unknown.

All actual anticoagulant rodenticides target the liver enzyme <u>V</u>itamin <u>K</u>ep<u>O</u>xide <u>R</u>eductase <u>C</u>omplex (VKORC). VKORC recycles vitamin K 2,3-epoxide to vitamin K hydroquinone, required to carboxylate several blood coagulation factors, such as II, VII, IX, and X ^10^. In particular, most anticoagulant molecules are antagonists of vitamin K. They bind to the VKORC protein 1 (VKORC1) at similar binding-pockets inhibiting vitamin K recycling, as suggested by the molecular structures of anticoagulants, and correlations between *vkorc1* gene mutations and resistance to anticoagulants ^5^. After several decades, only a few mutations correlating with rat anticoagulant resistance have been identified during many field surveys, in different countries and several laboratory tests. Most abundant mutations mapped in the *vkorc1* gene at its amino acid positions corresponding to L128Q, Y139C/S/F, L120Q and F63C (in the single letter amino acid code, first amino acid in wild type, last amino acid in the mutant). Rare mutations such as R33P, Y39N, A26T, and others, have been also described ^5, 11, 12, 14^. Similarly, in humans, where low concentrations of anticoagulants are clinically used in many diseases, only a few mutations on residues F55/G/Y, N80G and F83G have been implicated in binding to vitamin K by both computational and experimental methods^13^. Cystein residues conserved among species at C132/C135, were at the human VKORC1 catalytic site^13^.

Despite the above commented mutations, in a few cases, no mutations at the *vkorc1* gene ^15^, were detected in anticoagulant-resistance rats, suggesting that other genes could be also implicated. Among other possible genes, vertebrates also code for a *vkorc1* paralog gene, the *vkorc1*-*like 1* (*vkorc1l1*) ^16^. Although human VKORC1L showed different binding-sites and 2-100-fold lower binding-affinities to most anticoagulants compared to VKORC1 ^17, 18^, anticoagulant resistance, including that of rats, may implicate not only *vkorc1*, but also the *vkorc1l1* gene. On the other hand, anticoagulant resistance might as well implicate other still unknown genes, alterations of anticoagulant pharmacokinetics ^5, 12^ or detoxification mechanisms.

Enhancement of detoxifying enzymes, such as cytochrome P450 enzymes (CYPs), has been suggested by several authors to contribute to anticoagulant resistance, since their upregulation have been detected in anticoagulant resistant rats. For instance, bromadiolone-resistance rats homozygous for the Y139C VKORC1 mutation, upregulated *cyp1a2, cyp2c13, cyp2e1, cyp3a2* and *cyp3a3* among *3*9 other *cyp* genes which remained unchanged in comparison with their corresponding wild-type bromadiolone-susceptible rats^19^. Additionally, most of the modulated *cyp* genes were also upregulated after bromadiolone administration ^19^. All these results implicate not only *vkorc1/vkorc1l1 gene* mutations, but also *cyp* gene expression/activity in anticoagulant resistance^20^.

Although mammals have several enzymes for binding and degrading any foreign chemical compounds, the CYP heme-containing oxidative enzymes are the best studied. CYP enzymes, for instance, play a major role in metabolizing environmentally released compounds, binding a wide variety of chemicals ^21^. Furthermore, some CYPs even participate in normal metabolism, for instance, by generating a variety of hydroxycholesterols ^22^. Hundreds of *cyp*-like genes corresponding to the rat CYP1 to 4 enzyme superfamilies (rat genome database: http://rgd.mcw.edu/) and in other species including humans, have been putatively identified in their corresponding genomes (http://drnelson.uthsc.edu/cytochromeP450.html, http://drnelson.uthsc.edu/UNIGENE.RAT.html)^23^.

CYP enzymes exhibited species- and isomer-specific amino acid sequences, but they also show conserved carboxy-terminal segments and similarities in their tridimensional 3D structures. Therefore, CYPs specificity is mostly due to differences among their amino acid sequences rather than to their 3D structures. Specificities such as CYP1A2 binding of aromatic-flat molecules, CYP2C9 binding acid molecules, and CYP3A4 binding larger molecules are generally known. However, those oversimplifications may vary across species and/or even among sexes or different physiological situations within the same specie. Detoxification of one particular chemical compound even of different stereoisomers is more probable to implicate collaboration among several different CYPs and/or other hypothetical oxidative enzymes. For example, it is known that *S*-warfarin is detoxified by CYP2C9, while its stereo isomer *R*-warfarin, is detoxified by other CYPs ^24^. However, for any given compound, most of those complex detoxification pathways are largely unknown. In the particular case of rat CYPs potentially implicated in the resistance to different anticoagulants, it is also more probable that many CYPs/*cyps* and/or other still unknown enzymes/genes may be implicated.

As briefly reviewed above, competition among VKORC1 / VKORC1L1, CYPs, and different VKORC1 resistant mutants, may partially define the final effectivity of anticoagulant rodenticides. Due to the few reported anticoagulant computationally studies ^9^, the present work was focused on VKORC1, VKORC1L1, a selected group of CYP rat genes that changed their gene expression in anticoagulant-resistant rats ^19, 20^ and VKORC1 mutants. We explored their possible binding interactions with actual anticoagulant rodenticides and anticoagulant-similar molecular structures. We selected top-leads among anticoagulant-similar ligands with both VKORC1 binding-scores in the low nM range (high binding-affinities) and CYP high binding-scores (potential minimal detoxification). Those top-leads maintaining low binding-scores to the maximal number of known VKORC1 mutants (majority voting), were used to predict the top top-leads proposed for experimental validation.

## Materials and Methods

### Anticoagulant-like and vitamin K ligands

Anticoagulant-like ligands were obtained by similarity search in PubMed (https://pubchem.ncbi.nlm.nih.gov/) by providing anticoagulants in their SMILES formula. The anticoagulant provided were warfarin (PubChem ID 54678486), coumatetralyl (54678504), bromadiolone (54680085), difenacoum (54676884), brodifacoum (54680676), and flocoumafen (54698175). Vitamin K-like ligands were also obtained by similar search in PubMed. The corresponding downloaded sdf files were manually curated, merged into one sdf file, and duplicates eliminated using Open Babel vs 2.3.1. (http://openbabel.org/wiki/Windows_GUI) to render a final 653 ligand sdf file for docking.

### Tridimensional VKORC1 / VKORC1L1, CYP and VKORC1 mutant rat models

The amino acid sequences of rat VKORC1 / VKORC1L1, CYPs and VKORC1 mutants were translated from downloaded mRNA sequences present in the GenBank (http://www.ncbi.nlm.nih.gov/sites/entrez?db=nucleotide). The CYP enzymes selected for this work corresponded to those differentially expressed in anticoagulant resistant rats ^20, 25^ (Table 1).

**Table 1.** 3D modelling of VKORC1, VKORC1L1 and CYP rat enzymes.

| name | protein sequence | #aa | SWISS template | human template | CSI, % | RMSD, Å / atoms | ligand located in binding-pocket | R/S |
| --- | --- | --- | --- | --- | --- | --- | --- | --- |
| VKORC1 | NP_976080 | 161 | 4nv6 | VKORC1 | 100.0 | 1.00/161 | none described | ---- |
| VKORC1L1 | NM_203338 | 176 | 4nv6 | VKORC1 | 41.0 | 0.74/127 | none described | ---- |
| CYP1A2 | NM_012541 | 513 | 2hi4 | CYP1A2 | 100.0 | 1.00/513 | α-naphthoflavone | ND |
| CYP2C13 | NM_138514 | 490 | 5x23 | CYP2C9 | 66.8 | 1.98/411 | losartan | Co* |
| CYP2E1 | NM_031543 | 493 | 373z | CYP2E1 | 80.3 | 1.82/417 | pilocarpine | Ind |
| CYP3A2 | NM_153312 | 504 | 1tqn | CYP3A4 | 72.3 | 2.23/370 | progesterone | Ind |
| CYP3A3 | NM_013105 | 502 | 5a1r | CYP3A4 | 72.8 | 2.24/376 | progesterone | Ind |

The CYP enzymes corresponded to those differentially expressed in anticoagulant resistant rats<sup>20, 25</sup>. CSI, protein sequence identity with the template chosen by SWISS-MODEL. Ligand binding pocket, Binding pockets for VKORC1 were not defined while those of CYPs were given by the human models. #aa, number of amino acids. RMSD, Root Square Mean Differences, expressed in Å / number of common carbon atoms. The lower, the more similar. R/S, bromadiolone resistance/susceptible ratio of mRNA expression from microarray/RTqPCR data<sup>25</sup>. Co, constitutive expression. \*, only inducible in females. Ind, bromadiolone-inducible expression<sup>20</sup>. ND, not determined

Rat amino acid sequences were submitted to the SWISS-MODEL homology modelling (https://swissmodel.expasy.org/interactive) which automatically selected templates with the closest sequence identity (CSI), including the mutated amino acid sequences of VKORC1. For CYPs, the heme and ligand binding-pocket coordinates provided by the corresponding human CYP templates (**R**esearch **C**ollaboratory for **S**tructural **B**ioinformatics, RCSB, **P**rotein **D**ata **B**ank PDB), were copied to the corresponding files of the modeled rat pdb file. Modeled structures were visualized in PyMOL (https://www.pymol.org/). Tridimensional similarities were expressed in Angstroms Å / number of common carbon atoms, as estimated by calculating the Root Square Mean Differences (RMSD) of the alpha-carbons by 3D superposition of CYP1A2 with the rest of CYPs in the CCP4 Molecular Graphics program vs2.10.11 (http://www.ccp4.ac.uk/MG) (Table 1). Binding-pockets and α-helices were predicted (VKORC1 / VKORC1L1) or confirmed (CYPs) using seeSAR vs.10 (https://www.biosolveit.de/SeeSAR/).

### AutoDockVina virtual docking

The AutoDockVina program ^26^ included into the PyRx 0.9.8. package ^27^ (https://pyrx.sourceforge.io/) was used in e7 64-desk computers as described before ^22, 28, 29^, using grids for the whole VKORC1 / VKORC1L1 molecules (blind docking) or binding pockets centered around the heme molecules of CYPs. The *.sdf files were converted to *.pdbqt files after ffu energy minimization (Open Babel included into the PyRx package). The pose with the lowest binding-score (ΔG energy) of each *.out.pdbqt were converted to constant inhibition (Ki) in molar concentrations (M), using the formula Ki = exp([ΔG × 1000] / [R × T]) (R = 1.98 cal/mol, and T = 298 °C)^30^ and converted to nM. The predicted structures were visualized in PyRx and/or PyMOL.

### Virtual docking by the seeSAR package

The seeSAR vs.10 package (https://www.biosolveit.de/SeeSAR/)^31, 32^ including unfavorable interactions to reduce false positives ^33^ and employing different HYDE scoring functions compared to AutoDockVina ^26^, was chosen as an alternative docking methodology. To explore the CYP surfaces for binding, previously identified binding-pocket 3D coordinates were included into their rat pdb files (Table 1) and confirmed by seeSAR and/or PyMOL visualization before used for docking. The average between the lower and higher boundaries of the lowest binding-score prediction poses per ligand expressed in nM were selected for analysis.

## Results

### Characteristics of the 3D models of rat VKORC1 and VKORC1L1

Because of the absence of crystallization studies solving rat VKORC1/ VKORC1L1 3D structures, hypothetical models were obtained by automatic search of templates (Table 1 and Figure 1A). A preliminary study of the proposed 3D structure of rat VKORC1 fitted well with mapped mutations (Figure 1B), seeSAR predicted binding-pockets (Figure 1C) and our own preliminary computational best-pose of some anticoagulants (Figure S3,B).

**Figure 1.**
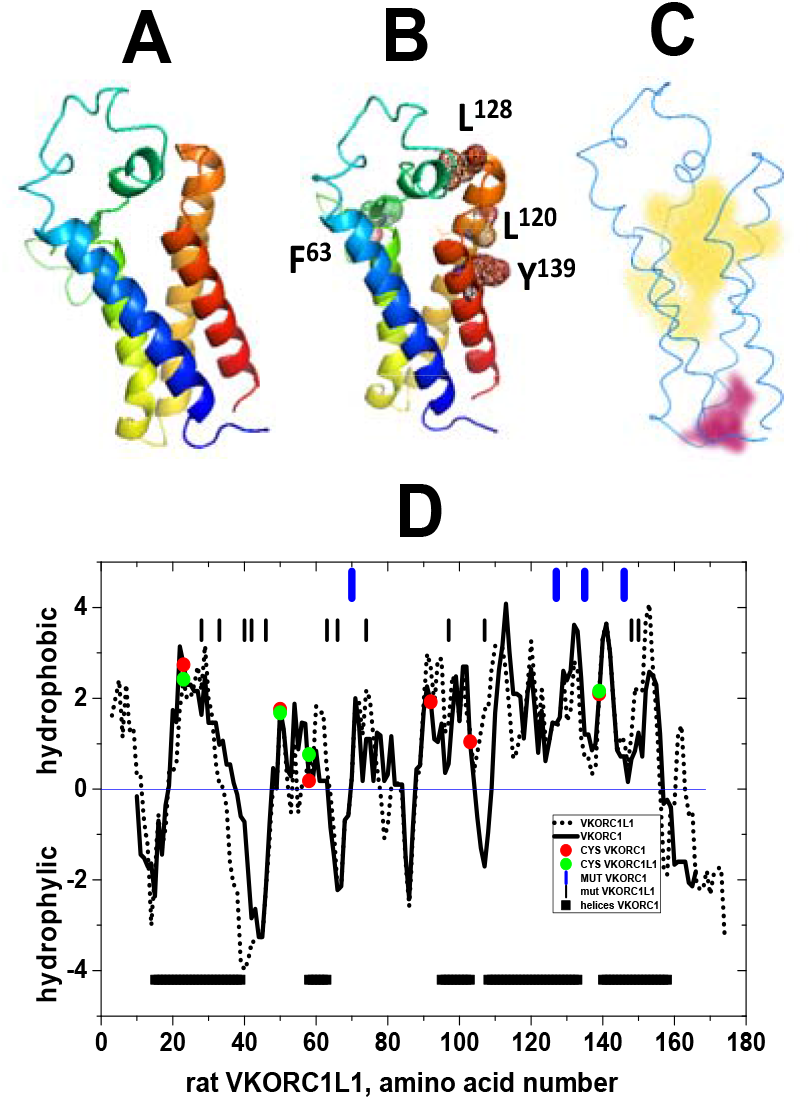
Properties and main mutant mappings of the amino acid sequence of rat VKORC1 and VKORC1L1. The amino acid sequences were from NP_976080 and NM_203338, for rat VKORC1 and VKORC1L1, respectively. Hydrophobic / hydrophilic plots were obtained by the Kyte & Doolittle values using Clone Manager vs9. Both structures were aligned by the location of 4 conserved cysteins, resulting in relative amino acid numbers corresponding to the 7 amino acid longer VKORC1L1 at its amino-terminal end. A) 3D predicted structures modeled from human VKORC1 4nv6 (**blue**, amino end; **red**, carboxy end). B) Most abundant amino acid mutations in VKORC1 correlating with rat anticoagulant resistance. C) Yellow and reddish backgrounds, binding-pockets predicted by seeSAR. **D) Black line and dotted lines**, VKORC1 and VKORC1L1 hydrophobic / hydrophilic profiles, respectively. **Red circles**, mapped cysteins in VKORC1. **Green circles**, mapped cysteins in VKORC1L1. **Blue vertical upper lines**, relative positions of the most abundant amino acid mutations in VKORC1 correlating with anticoagulant resistance. **Black vertical lines**, relative positions of minor amino acid mutations in VKORC1 correlating with anticoagulant resistance. **Horyzontal black rectangles**, relative positions of α-helices in VKORC1.

The modeled 3D structure of rat VKORC1 contains 5 short α-helices and 6 Cysteins (C), 4 of which are conserved among species. Cysteins C51-C132 form a disulphide bridge in humans^13^. Helices are highly hydrophobic at their carboxy-terminal segment (Figure 1D), containing the most abundant mutations correlating with rat anticoagulant resistance (Figure 1,B,D). VKORC1 and VKORC1L1 alignement suggested very similar hydrophobic/hydrophilic profiles (Figure 1D) and 3D structures (Table 1, 0.74 Å RMSD of VKORC1L1 compared to VKORC1), despite their differences in amino acid sequences (Table 1, 41 % CSI of VKORC1L1 compared to VKORC1). Thus, the sequence of VKORC1L1 although conserving four cystein positions and most of the hydrophobic profile compared to VKORC1, contained only 41 % identical amino acids, and included two additional amino acid segments at its amino and carboxy ends. Despite these differences in sequence, 3D superposition of the models predicted a low mean RMSD difference. The amino acid sequence differences were located mostly at the unstructured loops expanding residues ∼32-81 (VKORC1) and 40-87 (VKORC1L1) (not shown).The α-helices showed a complete alignement despite their amino acid sequence differences as analyzed by CCP4MG (not shown).

The seeSAR prediction of binding-pockets were similar but implicating different amino acids (Figure 1C and not shown). Computational predictions of the main binding-pocket by seeSAR correlated with some of the mapped mutations (Figure 1C, yellow binding-pocket). Other minor mutations mapped around either the main binding-pocket (Figure 1C) or the amino-terminal part of VKORC1, closer to the smaller binding-pocket (Figure 1C, reddish binding-pocket) (not shown).

Because compared to VKORC1, VKORC1L1 has been less studied, we included docking predictions of its binding to anticoagulant-likes to explore its possible participation.

### Comparison of anticoagulant-like ligand binding-scores between VKORC1 and VKORC1L1

Most of the binding-scores obtained by docking a subset of anticoagulant-like compounds to VKORC1 were ∼800-fold lower (higher binding-efficiency) than those for VKORC1L1 (Figure S1). Using arbitrary thresholds of <130 nM for VKORC1 and < 10^4^ nM for VKORC1L1, the leads could be distributed into 4 groups (Figure S1, blue horizontal and vertical lines). Among the groups showing the lowest binding-scores for VKORC1 there were 10 compounds and coumatetralyl (Figure S1, green circles in the upper left group) while those for VKORC1L1 were 3 compounds and bromadiolone (Figure S1, green circles to the bottom right group). There were also 8 anticoagulant-like ligands and flocoumafen which predicted similar binding-scores between VKORC1 and VKORC1L1 (Figure S1, yellow circles). Whether the association of these new anticoagulant-like ligands with some of their corresponding actual anticoagulants may be due to similar functional properties it is unknown.

VKORC1 and VKORC1L1 were further compared using a library of vitamin K-like compounds. In this case, binding-scores for VKORC1 were ∼10000-fold lower than those for VKORC1L1 (Figure S2). Leads to VKORC1 predicted 20 vitamin K-like ligands with lower binding-scores than vitamin K. As it was expected, all the vitamin K-like leads contained a similar coumarin ring. However, the ligands detected by seeSAR have additional 12-16 Carbon chain(s) with variations in the degree and location of additional hydroxylations and carboxylations.

Therefore, the above described results showed that most anticoagulant-like and vitamin K-like binding-scores were much lower (higher binding-efficacy) when bound to VKORC1 than to VKORC1L1, suggesting that targeting VKORC1 rather than VKORC1L1 may be more important for anticoagulation purposes. Therefore, further work was focused on VKORC1.

### Comparison of anticoagulant-like ligand binding-score profiles between AutoDockVina and seeSAR

Results of the anticoagulant-like ligands predicted to bind to VKORC1 by both AutoDockVina and seeSAR showed that there was correlation between the binding-score profiles of the best poses from both programs (not shown). However, the binding-scores predicted by AutoDockVina were 100-1000-fold lower than those for seeSAR (Figure S3). Furthermore, the number of ligands with lower binding-scores (leads) predicted by AutoDockVina were more numerous than those predicted by seeSAR. According to these profiles, anticoagulant binding predictions may be best obtained using AutoDockVina than seeSAR. Nevertheless it is possible that some additional chemical structures may be detected also by seeSAR.

Visual inspection of the lead-VKORC1 complexes predicted by either program showed that most of the leads mapped nearby the already mentioned resistance-linked mutations and binding-pockets (Figure 1B, C and Figure S3,A,B), confirming that the main binding site of VKORC1 may be correctly identified.

Because of all the results commented above, the AutoDockVina algorithm was selected for docking for the rest of analysis.

### Docking of actual anticoagulants and anticoagulant-like ligands to rat CYPs

The binding-scores after docking actual anticoagulants to rat CYPs correlated with their acute oral lethal dose fifty (LD^50^, expressed as mg of anticoagulant / Kg of body weight) (Figure 2). Results showed,

**Figure 2.**
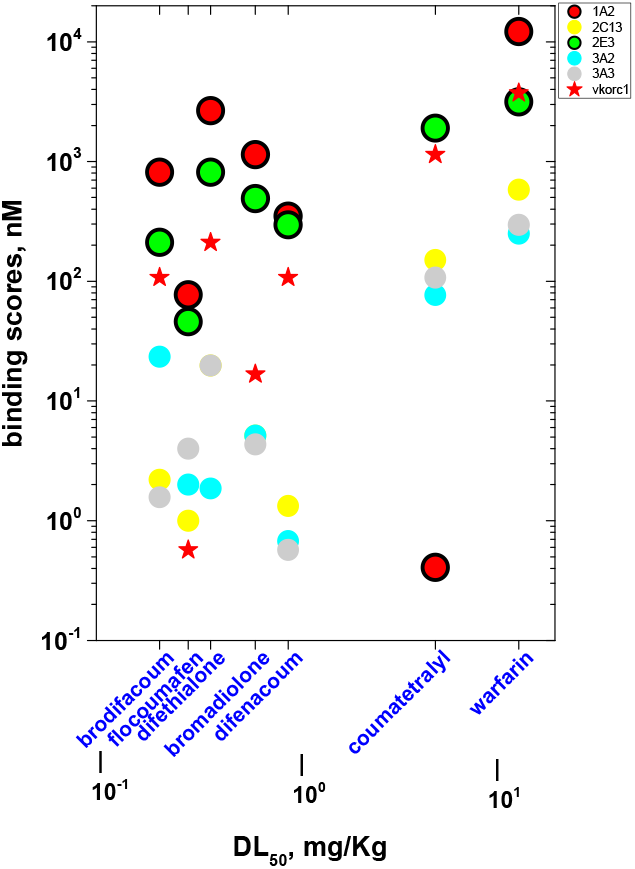
Binding-scores of actual anticoagulants docked to the selected rat CYPs. **Red circles**, CYP1A2. **Green circles** 2E3, **Red stars**, VKORC1. **Yellow circles**, CYP2C13. **Cyan circles**, 3A2. **Gray circles**, 3A3. **Bottom blue lettering**, actual anticoagulants ordered by their acute oral lethal dose fifty (LD50) in rats.

i. High binding-scores (∼10^3^-10^4^ nM) to CYP1A2 (Figure 2, red circles), and CYP2E3 (Figure 2, green circles) for most actual anticoagulants (except CYP1A2), compatible with low detoxification levels,
ii. Similar high binding-score profiles among CYP1A2, CYP2E3 and most VKORC1 (Figure 2, red stars),
iii. High binding-scores (∼10^2^-10^4^ nM) to all CYPs (except for CYP1A2), for those anticoagulants with higher LD^50^, such as warfarin and coumatetralyl,
iv. Low binding-scores (∼0.5-20 nM) to CYP2C13 (Figure 2, yellow circles), CYP3A2 (Figure 2, cyan circles) and CYP3A3 (Figure 2, gray circles) for those anticoagulants with lower LD^50^ such as difethialone, diphenadione, bromodifacoum, difenacoum, bromadiolone and flocoumafen.

These results obtained with actual anticoagulants of known *in vivo* rat LD^50^ activities, suggested that the high binding-scores to CYP1A2 and CYP2E3 may predict low detoxification levels for new anticoagulant-like ligands compatible with their possible higher anticoagulant activities. Therefore, the anticoagulant-like compounds were docked by AutoDockVina to rat VKORC1 and CYPs to identify top-leads with both lower VKORC1 and higher CYPs binding-scores, which in turn could correspond to higher anti-coagulation and lower detoxification capacities, favoring putative higher mortalities for rodenticide purposes.

Results comparing VKORC1 and CYPs showed binding-scores between 10^−2^ to 10^5^ and 10^−1^ to 10^5^ nM, respectively (Figure 3). To facilitate analysis of the results, the corresponding binding-scores of vitamin K (17 and 450 nM, respectively) were used as reference to arbitrarily define thresholds for VKORC1 (Figure 3, horyzontal hatched blue line) and CYPs (Figure 3, vertical hatched blue line). According to these arbitrary thresholds, ligands were divided in **A, B, C** and **D** groups (Figure 3).

**Figure 3.**
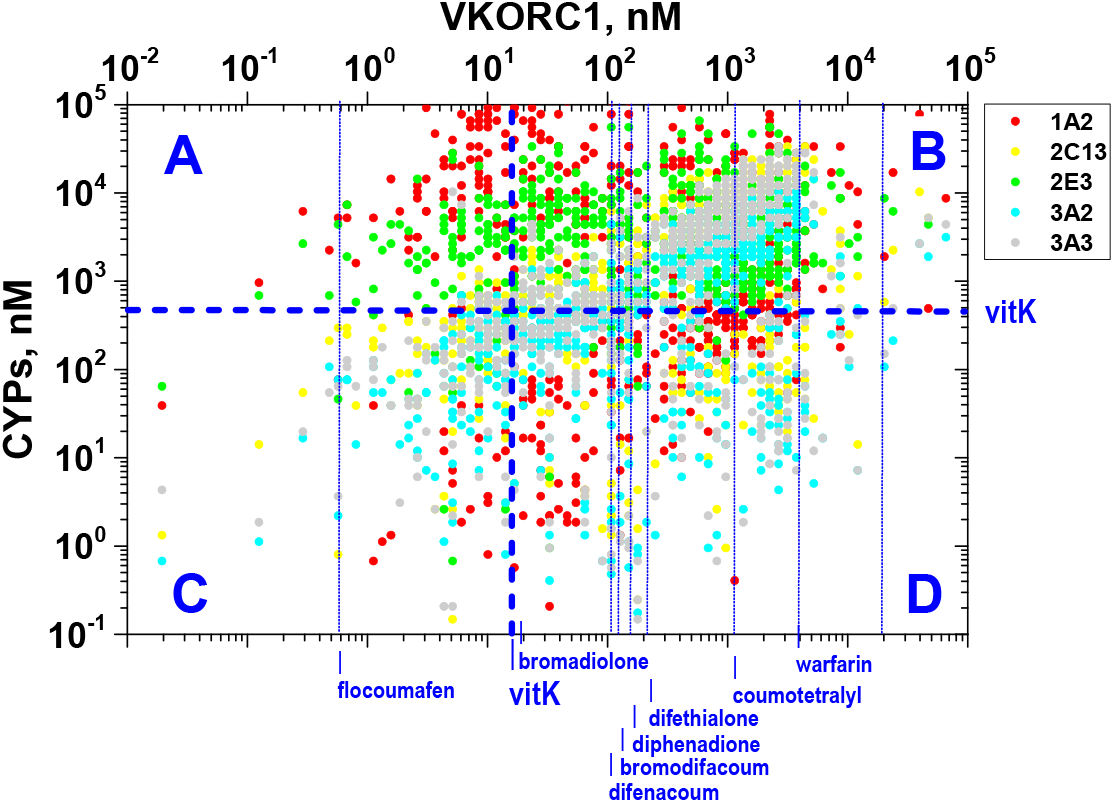
Comparison of binding-scores of VKORC1 and CYPs. The anticoagulant-like ligands were docked by AutoDockVina to VKORC1 and CYPs. Horyzontal and vertical blue dashed lines define the thresholds to separate groups A, B, C,D by the binding-scores defined by vitamin K-VKORC1 (Vertical) or CYPs mean (Horyzontal). **Blue A**, VKORC1 binding-scores <17 nM and >450 nM CYPs. **Blue B**, VKORC1 binding-scores >17 nM and >450 nM CYPs. **Blue C**, VKORC1 binding-scores <17 nM and <450 nM CYPs. **Blue D**, VKORC1 binding-scores >17 nM and <450 nM CYPs. **Blue lettering**, names of actual anticoagulants. **Red circles**, rat CYP1A2. **Yellow circles**, rat CYP2C13. **Green circles**, rat CYP2E1. **Cyan circles**, rat CYP3A2. **Gray circles**, rat CYP3A3.

Group **A** contained ligands with the lower binding-scores to VKORC1, suggesting their maximal anticoagulant activity, and higher CYP binding-scores suggesting lower detoxification levels. Most of the CYPs identified in group **A** were CYP1A2 and CYP3E1 (Figure 3**A**, red and green circles), thus providing adequate filters to select for those leads with minimal detoxification possibilities.

Group **C**, contained the lowest VKORC1 binding-scores, which may also be a source for alternative potent anticoagulants but with maximal detoxification levels. In contrast, groups **B** and **D**, contained ligands with VKORC1 high binding-scores which suggest that most of the ligands in this group would not improve actual anticoagulants.

Of the 112 VKORC1 leads of group **A**, 67 predicted high binding-scores for both CYP1A2 >1353 nM and 2E1 >3147 nM (thresholds of the corresponding binding-scores of vitamin K, respectively). To define top-leads, the rest of the CYPs were also computed and ranked by the total number of CYPs that were bound above their corresponding vitamin K binding-scores. Table 2 shows the resulting 41 top-leads defined by “ majority voting” among those VKORC1 leads which were bound by > 3 CYPs.

**Table 2.**
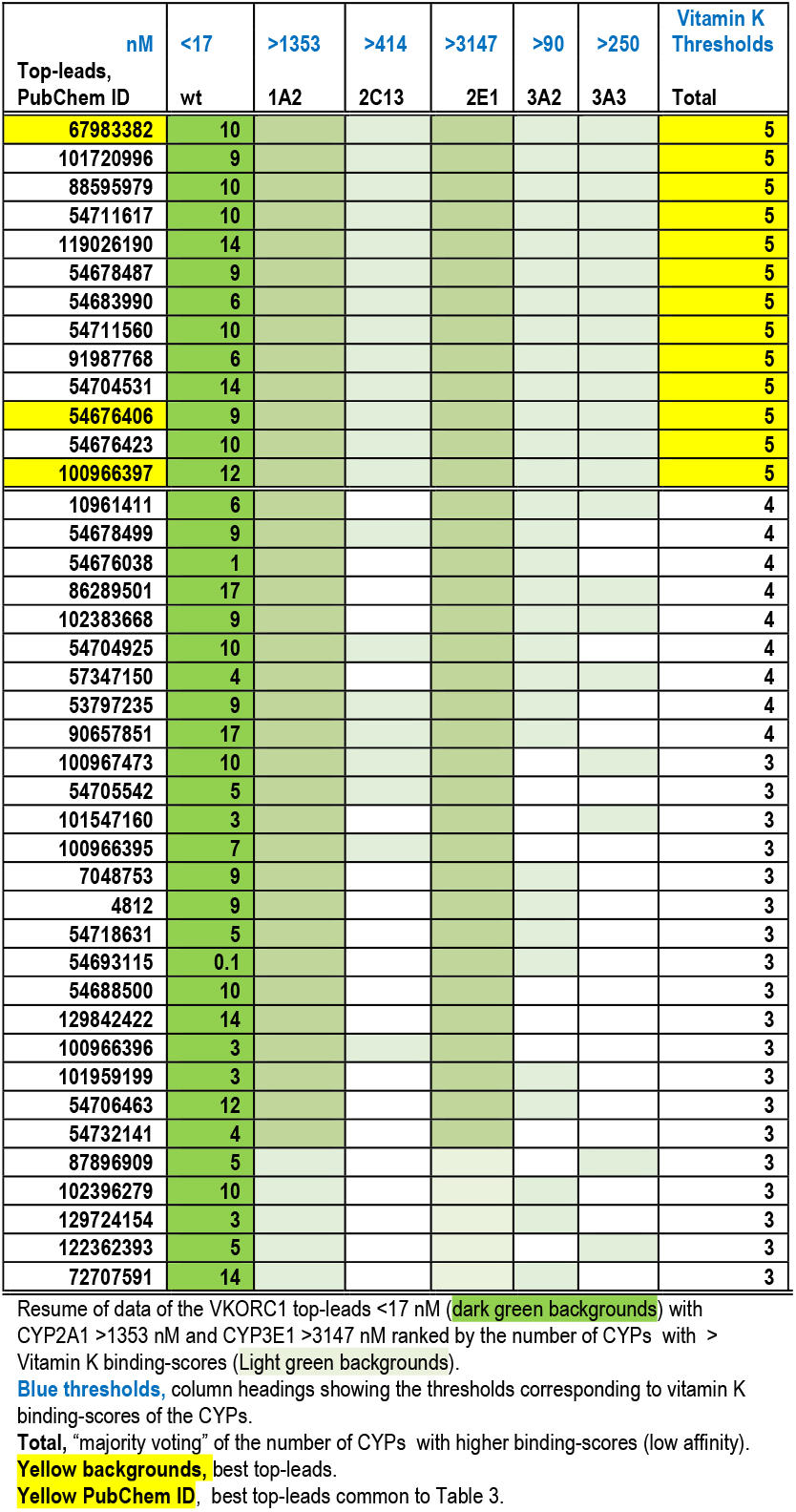
Anticoagulant-like VKORC1 top-leads and CYPs “ bottom-leads”.

Only 13 best top-leads were bound by the 5 CYPS studied (Table 2, Yellow Total). Most best top-leads were hydroxy-derived anticoagulant-like chemotypes of 4-hydroxy 1,2 benzopirone (hydroxycoumarin) which have an extra bencene ring linked to short carbon chains (chemotype **I**). This group included also Cl and Br derivatives (Figure 4). In addition, one of the best top-leads (ID 67983382) contained a naphthoquinone (1,4 naphtalenedione), a 2-ring structure with 2 oxo groups at positions 1 and 4 and a naphthalene ring linked to 19 hydrocarbon chains (chemotype **II**).

**Figure 4.**
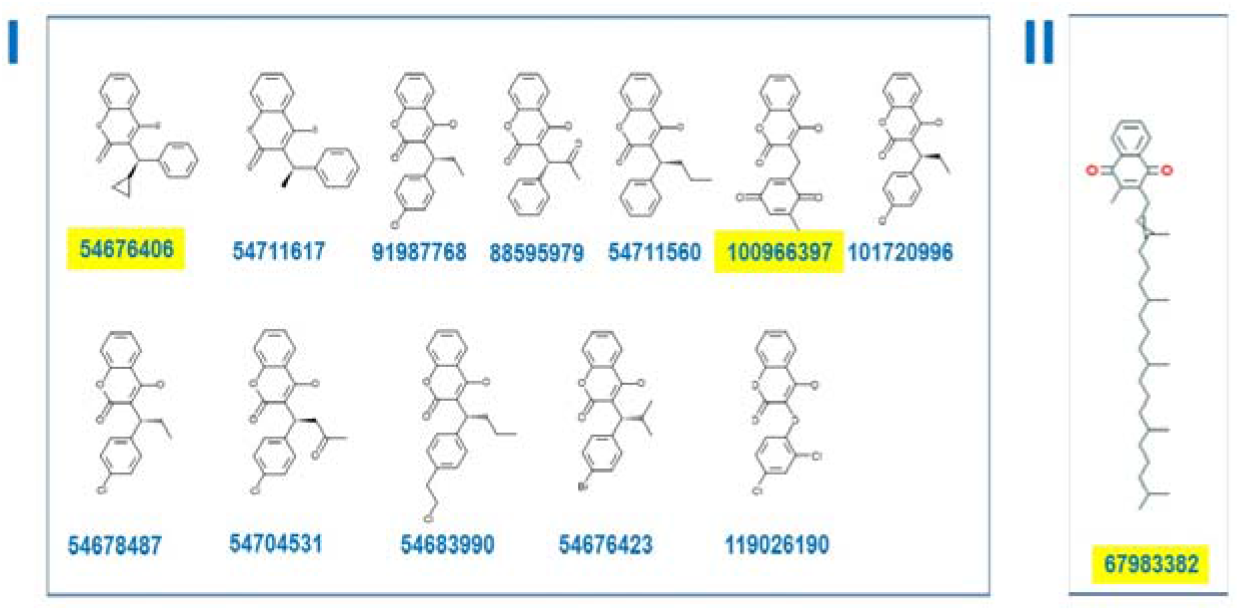
2D representation of top leads and CYP bottom leads. The best top-leads defined by the VKORC1 < 17 nM and weakly bound to CYPs (Table 2) were represented. **Blue bold numbers**, PubChem IDs. **Yellow PubChem ID**, best top-leads common to Table 3. **I**, hydroxycoumarin chemotypes. **II**, naphtoquinone chemotype.

The best top-lead compounds have low VKORC1, high CYP1A2 / 2E1 binding-scores, and high CYP2C13, CYP3A2 and CYP3A3 binding-scores, therefore they may be amongst the best candidates for anticoagulant rodenticide purposes (Table 2 and Figure 4).

**Table 3.**
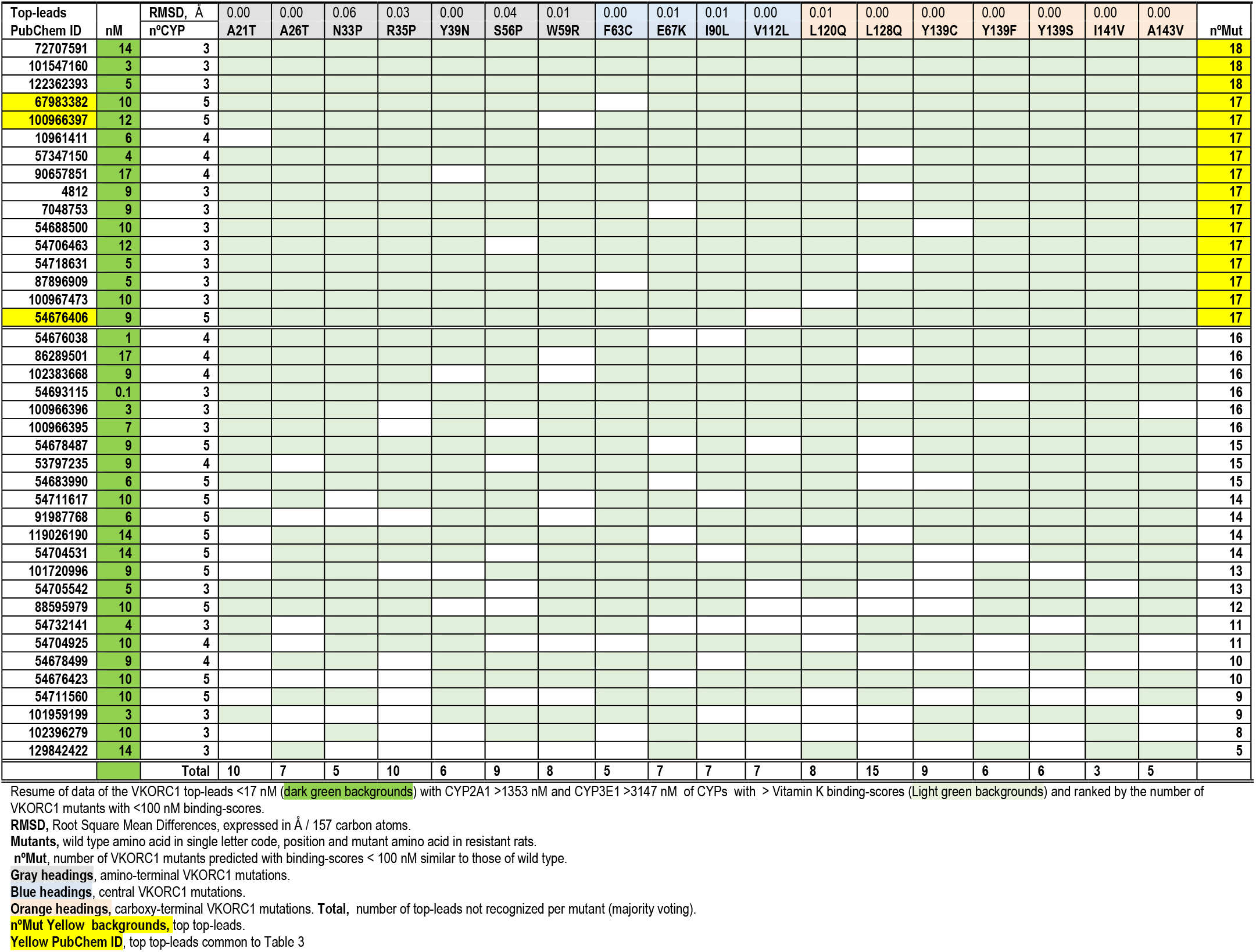
Anticoagulant-like VKORC1 top-leads, CYPs bottom-leads and mutant VKORC1 top-leads.

| Top-leads<br>PubChem ID | nM | RMSD, Å<br>n°CYP | 0.00<br>A21T | 0.00<br>A26T | 0.06<br>N33P | 0.03<br>R35P | 0.00<br>Y39N | 0.04<br>S56P | 0.01<br>W59R | 0.00<br>F63C | 0.01<br>E67K | 0.01<br>I90L | 0.00<br>V112L | 0.01<br>L120Q | 0.00<br>L128Q | 0.00<br>Y139C | 0.00<br>Y139F | 0.00<br>Y139S | 0.00<br>I141V | 0.00<br>A143V | n°Mut |
| --- | --- | --- | --- | --- | --- | --- | --- | --- | --- | --- | --- | --- | --- | --- | --- | --- | --- | --- | --- | --- | --- |
| 72707591 | 14 | 3 |  |  |  |  |  |  |  |  |  |  |  |  |  |  |  |  |  |  | 18 |
| 101547160 | 3 | 3 |  |  |  |  |  |  |  |  |  |  |  |  |  |  |  |  |  |  | 18 |
| 122362393 | 5 | 3 |  |  |  |  |  |  |  |  |  |  |  |  |  |  |  |  |  |  | 18 |
| 67983382 | 10 | 5 |  |  |  |  |  |  |  |  |  |  |  |  |  |  |  |  |  |  | 17 |
| 100966397 | 12 | 5 |  |  |  |  |  |  |  |  |  |  |  |  |  |  |  |  |  |  | 17 |
| 10961411 | 6 | 4 |  |  |  |  |  |  |  |  |  |  |  |  |  |  |  |  |  |  | 17 |
| 57347150 | 4 | 4 |  |  |  |  |  |  |  |  |  |  |  |  |  |  |  |  |  |  | 17 |
| 90657851 | 17 | 4 |  |  |  |  |  |  |  |  |  |  |  |  |  |  |  |  |  |  | 17 |
| 4812 | 9 | 3 |  |  |  |  |  |  |  |  |  |  |  |  |  |  |  |  |  |  | 17 |
| 7048753 | 9 | 3 |  |  |  |  |  |  |  |  |  |  |  |  |  |  |  |  |  |  | 17 |
| 54688500 | 10 | 3 |  |  |  |  |  |  |  |  |  |  |  |  |  |  |  |  |  |  | 17 |
| 54706463 | 12 | 3 |  |  |  |  |  |  |  |  |  |  |  |  |  |  |  |  |  |  | 17 |
| 54718631 | 5 | 3 |  |  |  |  |  |  |  |  |  |  |  |  |  |  |  |  |  |  | 17 |
| 87896909 | 5 | 3 |  |  |  |  |  |  |  |  |  |  |  |  |  |  |  |  |  |  | 17 |
| 100967473 | 10 | 3 |  |  |  |  |  |  |  |  |  |  |  |  |  |  |  |  |  |  | 17 |
| 54676406 | 9 | 5 |  |  |  |  |  |  |  |  |  |  |  |  |  |  |  |  |  |  | 17 |
| 54676038 | 1 | 4 |  |  |  |  |  |  |  |  |  |  |  |  |  |  |  |  |  |  | 16 |
| 86289501 | 17 | 4 |  |  |  |  |  |  |  |  |  |  |  |  |  |  |  |  |  |  | 16 |
| 102383668 | 9 | 4 |  |  |  |  |  |  |  |  |  |  |  |  |  |  |  |  |  |  | 16 |
| 54693115 | 0.1 | 3 |  |  |  |  |  |  |  |  |  |  |  |  |  |  |  |  |  |  | 16 |
| 100966396 | 3 | 3 |  |  |  |  |  |  |  |  |  |  |  |  |  |  |  |  |  |  | 16 |
| 100966395 | 7 | 3 |  |  |  |  |  |  |  |  |  |  |  |  |  |  |  |  |  |  | 16 |
| 54678487 | 9 | 5 |  |  |  |  |  |  |  |  |  |  |  |  |  |  |  |  |  |  | 15 |
| 53797235 | 9 | 4 |  |  |  |  |  |  |  |  |  |  |  |  |  |  |  |  |  |  | 15 |
| 54683990 | 6 | 5 |  |  |  |  |  |  |  |  |  |  |  |  |  |  |  |  |  |  | 15 |
| 54711617 | 10 | 5 |  |  |  |  |  |  |  |  |  |  |  |  |  |  |  |  |  |  | 14 |
| 91987768 | 6 | 5 |  |  |  |  |  |  |  |  |  |  |  |  |  |  |  |  |  |  | 14 |
| 119026190 | 14 | 5 |  |  |  |  |  |  |  |  |  |  |  |  |  |  |  |  |  |  | 14 |
| 54704531 | 14 | 5 |  |  |  |  |  |  |  |  |  |  |  |  |  |  |  |  |  |  | 14 |
| 101720996 | 9 | 5 |  |  |  |  |  |  |  |  |  |  |  |  |  |  |  |  |  |  | 13 |
| 54705542 | 5 | 3 |  |  |  |  |  |  |  |  |  |  |  |  |  |  |  |  |  |  | 13 |
| 88595979 | 10 | 5 |  |  |  |  |  |  |  |  |  |  |  |  |  |  |  |  |  |  | 12 |
| 54732141 | 4 | 3 |  |  |  |  |  |  |  |  |  |  |  |  |  |  |  |  |  |  | 11 |
| 54704925 | 10 | 4 |  |  |  |  |  |  |  |  |  |  |  |  |  |  |  |  |  |  | 11 |
| 54678499 | 9 | 4 |  |  |  |  |  |  |  |  |  |  |  |  |  |  |  |  |  |  | 10 |
| 54676423 | 10 | 5 |  |  |  |  |  |  |  |  |  |  |  |  |  |  |  |  |  |  | 10 |
| 54711560 | 10 | 5 |  |  |  |  |  |  |  |  |  |  |  |  |  |  |  |  |  |  | 9 |
| 101959199 | 3 | 3 |  |  |  |  |  |  |  |  |  |  |  |  |  |  |  |  |  |  | 9 |
| 102396279 | 10 | 3 |  |  |  |  |  |  |  |  |  |  |  |  |  |  |  |  |  |  | 8 |
| 129842422 | 14 | 3 |  |  |  |  |  |  |  |  |  |  |  |  |  |  |  |  |  |  | 5 |
| Total |  |  | 10 | 7 | 5 | 10 | 6 | 9 | 8 | 5 | 7 | 7 | 7 | 8 | 15 | 9 | 6 | 6 | 3 | 5 |  |

Resume of data of the VKORC1 top-leads <17 nM (dark green backgrounds) with CYP2A1 >1353 nM and CYP3E1 >3147 nM of CYPs with > Vitamin K binding-scores (Light green backgrounds) and ranked by the number of VKORC1 mutants with <100 nM binding-scores.
RMSD, Root Square Mean Differences, expressed in Å / 157 carbon atoms.
Mutants, wild type amino acid in single letter code, position and mutant amino acid in resistant rats.
n°Mut, number of VKORC1 mutants predicted with binding-scores < 100 nM similar to those of wild type.
Gray headings, amino-terminal VKORC1 mutations.
Blue headings, central VKORC1 mutations.
Orange headings, carboxy-terminal VKORC1 mutations. Total, number of top-leads not recognized per mutant (majority voting).
n°Mut Yellow backgrounds, top top-leads.
Yellow PubChem ID, top top-leads common to Table 3

### Docking anticoagulant-like top-leads to VKORC1 mutants

Since most actual anticoagulants and top-leads targeted VKORC1 similar binding-pockets than vitamin K (Figure S1), most resistant VKORC1 mutations had to compete for survival with vitamin K binding. The relatively few resistant VKORC1 mutations found in years of rodenticide use underline the biological difficulties of generating such mutations. Therefore, an additional screening criteria for novel anticoagulants would be to select those with the additional criteria of maintaining low binding-scores not only to wild-type VKORC1 but also to its mutants. To this end the 41 top-leads of Table 2 were screened also for binding to VKORC1 mutants.

Results showed that 15 top-leads (top top-leads) bound with binding-scores < 100 nM to >17-18 VKORC1 known mutants (Table 3). These top top-leads contained 6 top-leads with the highest binding-scores to 4-5 CYPs, and wild-type VKORC1 with binding-scores between 3-17 nM. There were no apparent correlations between binding-scores and mutations in the amino-, central, or carboxy-segments of VKORC1, nor with possibly altered conformations upon mutation (RMSDs were very similar among all mutants)(Table 3, head).

About half of the top top-leads were hydroxycoumarin-like chemotypes (Figure 5, chemotype **I**) some of them showing one hydroxy group in bencene rings (IDs 72707591, 122362393) or a second hydrocarbon chain (ID 122362393). The rest were naphthoquinone-like compounds (Figure 5, chemotype **II**) with extra oxygens, methyl positions or stereoisomers and/or different double bonds.

**Figure 5.**
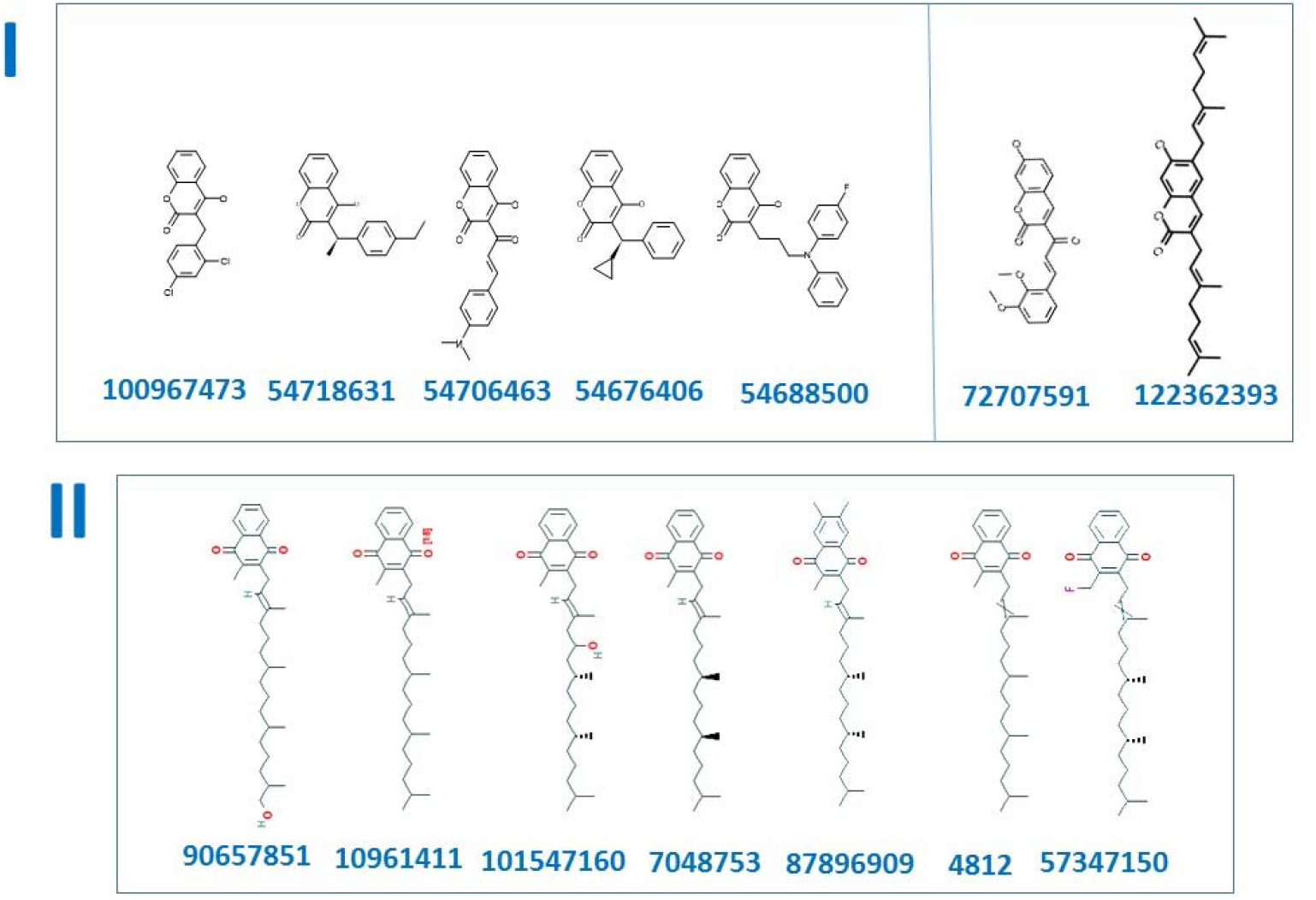
2D representation of VKORC1 top-leads, CYP bottom-leads and VKORC1 mutant top-leads. The top-leads defined by the VKORC1 < 17 nM, weakly bound to 5 CYPs (Table 2) and binding to >17 VKORC1 mutants (Table 3) were represented. **Blue bold numbers**, PubChem IDs. The top-leads common to those compounds listed on Figure 3 (yellow backgrounds) were not repeated here. **I**, hydroxycoumarin-like chemotypes. **II**, naphtoquinone-like chemotypes.

Chemotypes **I** and **II** may be amongst the best candidates for new anticoagulant rodenticide purposes because of their putative higher anticoagulant and lower detoxification activities while maintaining their binding levels to most of the VKORC1 mutants correlating with anticoagulant-resistance (Table 3 and Figure 5).

## Discussion

We describe here an approach to suggest novel rodenticides by computational targeting a combination of selected host enzymes. The results predicted a list of candidates with optimal ligand properties to test for possible anticoagulant rodenticide activities.

Our approach was based on computational binding screening of hundreds of anticoagulant-like ligands to rat VKORC1, CYPs and mutant-VKORC1 enzymes. Those targets were selected after discarding VKORC1L1 as a possible alternative or complement to VKORC1. Additional work was focused on VKORC1 because it predicted several order of magnitude lower binding-scores (higher binding-affinities) to several anticoagulant-like candidates, confirming previous results obtained by other authors^18^. Among the possible detoxification enzyme alternatives, CYPs appeared as the best predictors for our present purposes because they are the most important detoxificants in animals, including rats and only some rat CYPs have shown *in vivo* experimental evidences of correlations between anticoagulant resistance and changes in gene expression and/or anticoagulant administration and gene regulation.

Our initial hypothesis was that more potent anticoagulant-like rodenticide candidates could be predicted for experimental validation among those leads (top-leads) having not only maximal VKORC1 binding for best anticoagulant activity (similar to those reported before ^9^), but also minimal detoxification that would theoretically predict to expand their physiological time as poison.

Furthermore, top top-leads which were additionally predicted to bind to known anticoagulant resistant VKORC1 mutants, could minimize any future appearance of those mutations and help to downsize experimental validation efforts by adding an additional selective criteria.

Our results ranked some optimal candidates among a wide list of possibilities, since many variables affect *in silico* predictions. For instance, different results could had been predicted depending on the initial library of ligands screened, for instance although an anticoagulant-like library was used, many other larger libraries could have been targeted such as those containing natural compounds or synthetic chemicals. The algorithm program used for screening whether be AutoDockVina or seeSAR or many other, would be another important variable. The arbitrary thresholds used to downsize the resulting docked lists also influenced the outcome, thus besides the thresholds based on Vitamin K binding-scores used, many other arbitrary alternatives were possible. The criteria of majority voting that was used to rank top-leads, gave the same weight to each of the VKORC1, CYPs and mutant predictions, but other weights could have been chosen. Finally, the order of filtering VKORC1-CYPs-mutants rather than CYPs-VKORC1-mutants or VKORC1-mutants-CYPs would alter also the rankings. The final list of top top-leads proposed in Table 3 for experimental evaluation will be influenced and will vary for each of the different analysis strategies mentioned above. Therefore, different optimal candidates could have been computationally predicted for experimental validation.

Our initial hypothesis been that anticoagulant rodenticides may be improved among those showing both higher binding-affinity to VKORC1 together with low binding-affinity to CYPs, was successful in the sense that we could find top-lead candidates theoretically improving some of the properties of actual anticoagulants. However, the particular predicted properties such as anticoagulant and detoxification have not been yet experimentally defined in terms of binding-scores. Also, many other *in vivo* variables have not been taken into account to possible restrict *in silico* predictions. For instance, a desirable criteria for anticoagulants is to delay their induced mortality to avoid rat “ learning from rapid killing”, however such additional criteria has difficulties to be applied in a molecular quantifiable manner. Despite those difficulties, some inverse correlation have been found between the ranking of *in vivo* potency of actual anticoagulant rodenticides and their VKORC1 and CYPs binding-scores or the number of rat CYPs which could bind them (Figure 2). These results suggest that the strategy followed here may partially predict some real anticoagulant properties. Perhaps similar screening among natural compounds ^22, 28, 29^ or synthetic chemical libraries rather than anticoagulant-likes, could reveal novel chemotypes with the appropriated combination of lower and higher binding-scores to VKORC1, VKORC1 mutants and CYPs, respectively.

An additional concern is the possibility that novel compounds after being used as rodenticides would induce new VKORC1 resistant mutations. In other words, would exposed rats eventually become also genetically resistant?. Because the binding to VKORC1 of actual anticoagulants and most of the anticoagulant-like top-leads described here, mapped into similar vitamin K-binding-pockets, it seems likely that most other mutations would compete also with Vitamin K binding. Vitamin K binding is required for rats for successful coagulation and survival. Even those mutations mapping in other, possible allosteric VKORC1 binding-pockets would compete with normal coagulation and survival. These may explain the world-wide difficulties of rats to develop VKORC1 resistance to anticoagulants, as evidenced by the relatively low number of mutations detected after decades of use (i.e., a maximal of ∼ 20 mutants of which only 4-6 were highly prevalent). Therefore, although mutations against the new ligands could appear, history argues against many possibilities for other novel resistance mutations on the rat *vkorc1* gene that could simultaneously maintain their vitamin K binding.

Therefore, screening among VKORC1/CYPs to known VKORC1 mutants, may offer a way to improve their chances as practical anticoagulants. However, only time and experimental evidence would validate these assumptions.

Although detoxification of any compounds can be studied by *in vivo* and *in vitro* assays on invertebrates and fish, *in silico* predictions have been previously used to reduce the number of such tests ^34-37^. The approach used here is similar to other *in silico* prediction models for metabolism and toxicity, such as those provided by numerous ADMET software. Those programs could be applied to the anticoagulant rodenticide chemical structures. Nevertheless, such models generally use human CYPs and have not included rat CYPs, nor those rat CYPs correlating with rodenticide resistance. On the other hand, few reports do exist on such *in silico* predictions on anticoagulant rodenticides ^38^.

Experimental pre-validation studies of the top top-leads described here may include recombinant VKORC1 for *in vitro* solid-phase binding assays, CYP toxicity by *in vitro* assays employing cell lines derived from rat liver ^39^, etc. Experimental evidences from those or any other *in vitro* assays may provide indications as to whether any of these newly described molecules have increased possibilities to be relevant for anticoagulant rodenticide purposes, before the *in vivo* definitive but costly assays are carried out with a reduced number. In addition to their possible anticoagulant rodenticide activities, any of these new molecules may also be used as tools to continue the study of possible VKORC1 / CYP relationships.

## SUPPORTING INFORMATION

**Figure S1.**
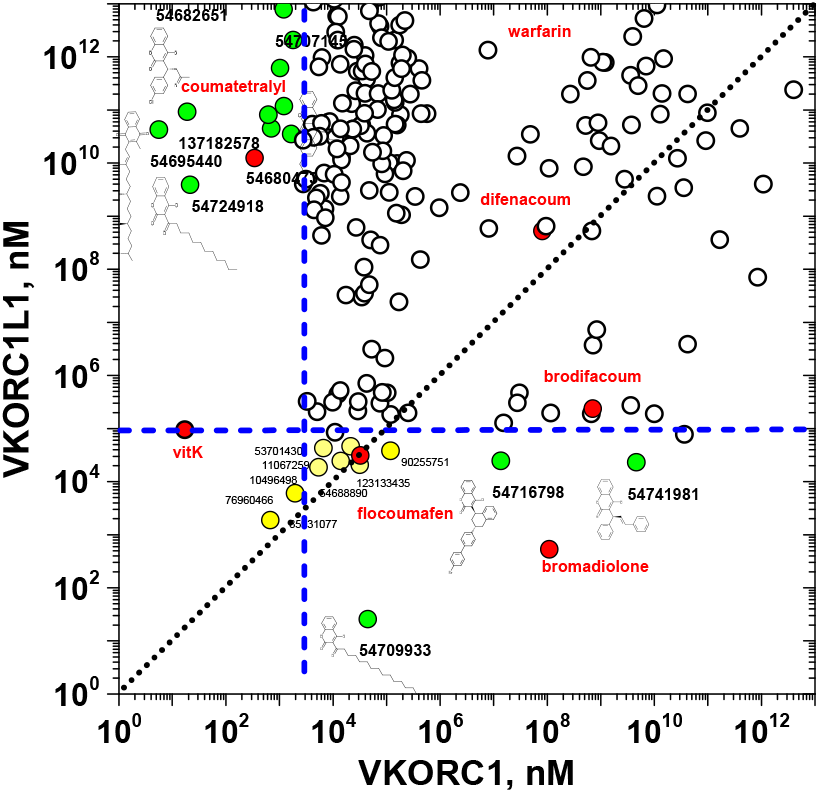
Comparison of VKORC1 and VKORC1L1 seeSAR binding-scores. Chemical formulas are included in some examples. **Black point line**, same binding-scores. **Blue vertical and horyzontal lines**, binding-score thresholds were arbitrarily defined for VKORC1 (130 nM) and VKORC1L1 (100000 nM). **Red circles and text**, actual anticoagulants and vitamin K. **Green circles**, ligands with binding-scores <thresholds. **Yellow circles**, ligands with similar binding-scores to VKORC1 and VKORC1L1. **Numbers**, PubChem IDs.

**Figure S2.**
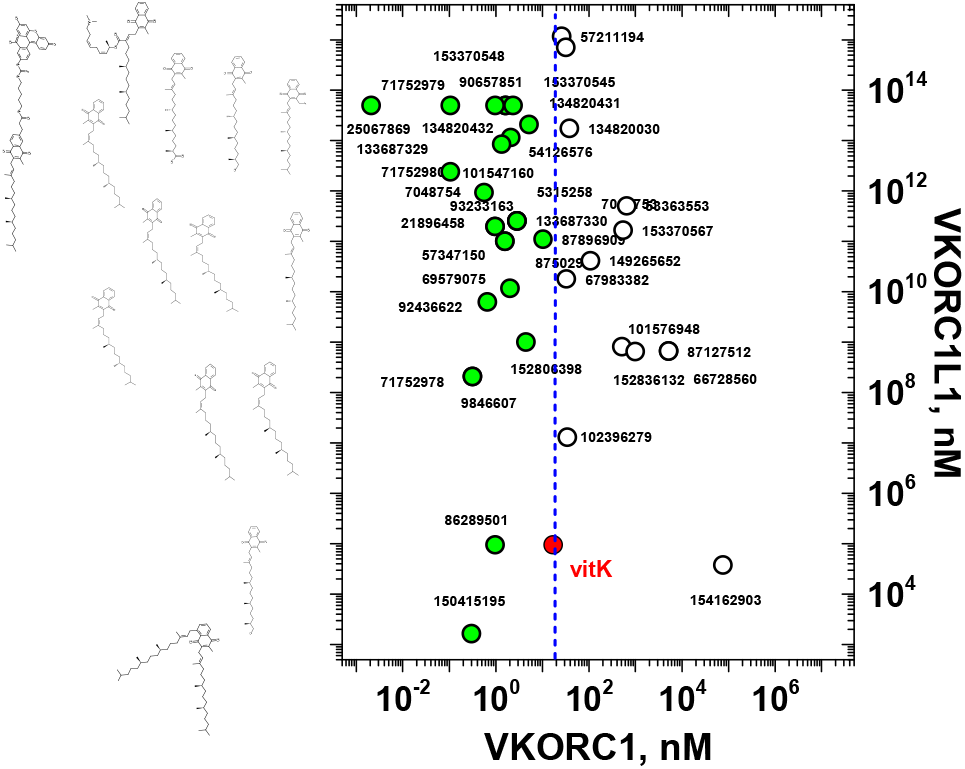
Comparison of VKORC1 and VKORC1L1 seeSAR binding-scores to vitamin K-like compounds. The corresponding formulas to the leads with binding-scores < 17 nM are shown to the left. **Blue vertical line**, threshold for leads to VKORC1 defined by the vitamin K binding-score (17 nM). **Numbers**, PubChem IDs.

**Figure S3.**
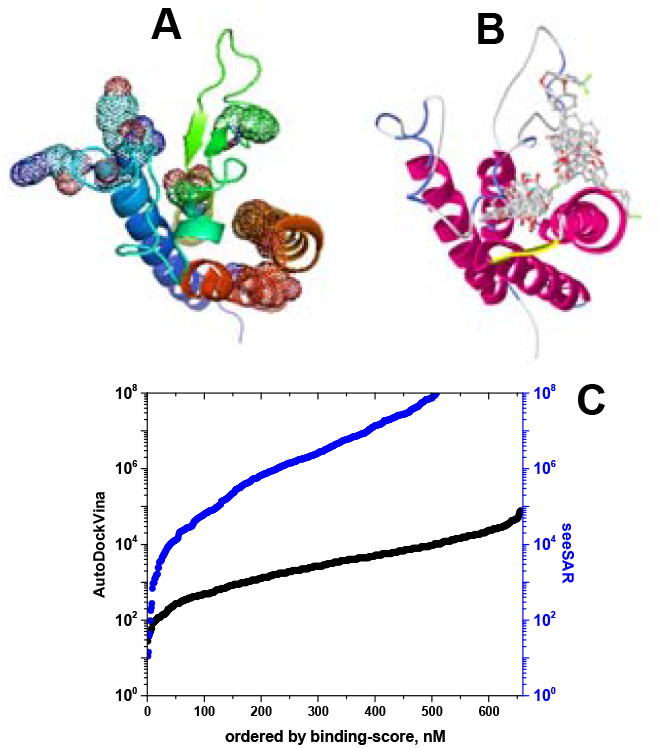
Comparison of VKORC1 binding-scores to anticoagulant-like ligands by seeSAR and AutoDock Vina. **A**. Mapping of the main anticoagulant resistance mutants in dotted spheres drawn in PyMOL. **B**, Mapping of the binding sites of several leads after AutoDockVina docking drawn in PyRx (similar maps were obtained by seeSAR docking). **C**. Comparison of binding-scores between AutoDockVina and seeSAR algorithms. The AutodockVina data were repeated in two different computers and the mean values calculated and converted to nM. The seeSAR data were calculated as the mean between lower and higher boundary estimations of binding-scores in nM. **Blue line**, seeSAR. **Black line**, AutoDockVina.

## Authors’ contributions

AB, collaborated in the docking, bibliography and discussions. JMN, coordinated the work and collaborated in the discussions. JC performed and analyzed the dockings, and drafted the manuscript. All authors read and approved the manuscript.

## Acknowledgements

Thanks are due to Dra Ana Valdehita for her help to obtain additional seeSAR licenses.

